# Cell type specific profiling of alternative translation identifies novel protein isoforms in the mouse brain

**DOI:** 10.1101/324236

**Authors:** Darshan Sapkota, Allison M. Lake, Wei Yang, Chengran Yang, Hendrik Wesseling, Amanda Guise, Ceren Uncu, Jasbir S. Dalal, Andrew Kraft, Jin-Moo Lee, Mark S. Sands, Judith A. Steen, Joseph D. Dougherty

## Abstract

Translation canonically begins at a single AUG and terminates at the stop codon, generating one protein species per transcript. However, some transcripts may use alternative initiation sites or sustain translation past their stop codon, generating multiple protein isoforms. Through other mechanisms such as alternative splicing, both neurons and glia exhibit remarkable transcriptional diversity, and these other forms of post-transcriptional regulation are impacted by neural activity and disease. Here, using ribosome footprinting, we demonstrate that alternative translation is likewise abundant in the central nervous system and modulated by stimulation and disease. First, in neuron/glia mixed cultures we identify hundreds of transcripts with alternative initiation sites and confirm the protein isoforms corresponding to a subset of these sites by mass spectrometry. Many of them modulate their alternative initiation in response to KCl stimulation, indicating activity-dependent regulation of this phenomenon. Next, we detect several transcripts undergoing stop codon readthrough thus generating novel C-terminally-extended protein isoforms *in vitro*. Further, by coupling Translating Ribosome Affinity Purification to ribosome footprinting to enable cell-type specific analysis *in vivo*, we find that several of both neuronal and astrocytic transcripts undergo readthrough in the mouse brain. Functional analyses of one of these transcripts, *Aqp4*, reveals readthrough confers perivascular localization, indicating readthrough can be a conserved mechanism to modulate protein function. Finally, we show that AQP4 readthrough is disrupted in multiple gliotic disease models. Our study demonstrates the extensive and regulated use of alternative translational events in the brain and indicates that some of these events alter key protein properties.

## Significance

Here we examine the extent and function of unusual protein variants resulting from alternative modes of mRNA translation in the mouse brain. We unexpectedly find that hundreds of neural transcripts use non-canonical translation initiation sites generating N-terminal protein variants, and dozens undergo stop codon readthrough generating C-terminally-extended protein variants. Further, we find that many transcripts modulate their use of non-canonical initiation in response to neuronal stimulation and at least one transcript modules its readthrough in response to diseases or conditions involving gliosis. Thus, our results demonstrate a novel use of alternative translation in homeostatic functions as well as in disease pathogenesis in the brain.

## Introduction

“One gene, one protein” has long been an obsolete axiom. Cells use alternative splicing, a process in which a single gene generates multiple transcripts, each of which may be translated into a protein isoform^1^. RNA-Seq studies on human cells have shown that 92–97% of multi-exon genes use alternative splicing^2,3^. To further expand proteomic landscapes, cells can also utilize alternative initiation and termination to generate multiple protein isoforms per transcript. While alternative splicing clearly is modulated by both neural activity as well as disease states^4–7^, alternative translation has not been systematically studied in the brain.

Alternative translation occurs from differences in initiation or termination on a given transcript. In conventional eukaryotic initiation, a 43S preinitiation complex scans the 5’ untranslated region (UTR) until it encounters an AUG at the annotated translation initiation site (aTIS) where it is joined by the 60S ribosomal subunit to form an 80S initiation complex^8^. However, an 80S complex may assemble at an upstream translation initiation site (uTIS) if the 5’UTR has an optimal AUG (or near-cognate NUG) or at a downstream translation initiation site (dTIS) if the aTIS is suboptimal^9^. Similarly, in conventional termination, the 80S complex concludes translation at the first in-frame stop codon. For some transcripts, however, it may read past that stop codon at some frequency and conclude translation at a second stop codon in the 3’UTR^10^. As expected, alternative initiation may generate N-terminal protein variants when in frame or entirely different protein species when out of frame, whereas stop codon readthrough can give rise to a C-terminally extended protein variants (Fig 1A). Thus, “one transcript, one protein” also is not a universally true axiom.

**Figure 1.**
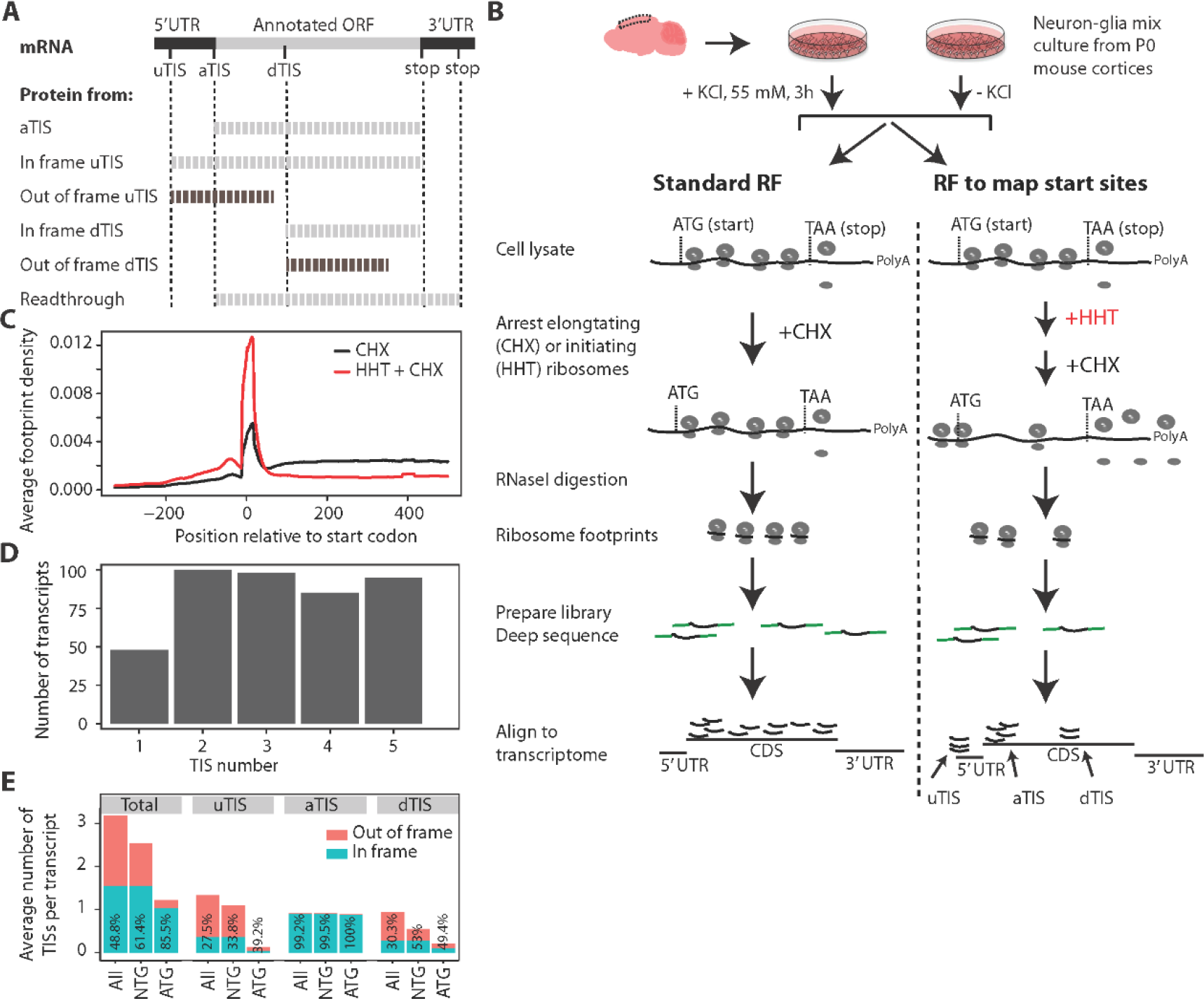
RF of homoharringtonin-treated neuron/glia culture reveals translation initiation sites. **A)** uTIS, aTIS and dTIS and the corresponding protein products are depicted. uTIS and dTIS give rise to N terminal variants of the protein when in frame, but code for completely new polypetide sequences when out of frame. Stop codon readthrough that generates a C-terminal extension is also depicted. In this study, Figures 1 and 2 concern TISs and Figures 3, 4 and 5 concern readthrough. **B)** Experimental workflow for *in vitro* study. Mixed neurons and glia from P0 mice were cultured for 7 days in vitro, exposed to KCl or no KCl, and subjected to RF and TIS-mapping RF as shown. **C)** Average ribosomal density across transcripts shows a run-off of elongating ribosomes from the proximal 300 nt region of the coding sequence in the HHT-treated sample as compared to the no HHT sample. Plot from the KCl-untreated cultures is shown; KCl-treated cultures gave the same density distributions. **D)** The number of high fidelity TISs in the 426 most robustly expressed transcripts. Most of the transcripts showed >1 TIS. Only top-five TISs were called. **E)** Codon composition and frame status of TISs across transcripts. TIS codons are shown as ATG or a cognate thereof (NTG). Numbers inside the bars indicate the percentages of in-frame TISs. RF, ribosome footprinting; HHT, homoharringtonin; TIS, translation initiation site; u/a/dTIS, upstream/annotated/downstream TIS.

In tissues other than central nervous system (CNS), genomic and proteomic experiments have revealed a widespread use of alternative TISs^11–17^ and readthrough^18,19^. In CNS cells, single gene studies have identified alternative open reading frames (ORFs) for individual transcripts and demonstrated their functional roles in homeostasis and disease^20–28^. However, there have been no systematic surveys of alternative translation across the brain or assessment of its variation across neurons and astrocytes. Here we aimed to define the alternative translation events *in vitro* and *in vivo* in the CNS. Furthermore, to more deeply understand their propensity for regulation, we tested the hypothesis these events are modulated by neural stimulation and disease states.

We screened for robust uTISs and dTISs in mixed neuron-glia cultures genome-wide using ribosome footprinting (RF), validated a subset of these with liquid chromatography tandem mass spectrometry (LC-MS/MS), and then determined how their use is impacted by KCl depolarization. We also identified transcripts evidencing stop codon read-through in cultures. We then coupled RF to translating ribosome affinity purification (TRAP) to determine whether neurons and astrocytes use readthrough *in vivo*. We found that several dozen transcripts, both cell type-specific and non-specific, undergo measurable readthrough and that one event, readthrough of *Aqp4* alters protein localization in the mouse brain and is dysregulated in models of neurological disease.

## Results

### RF of homoharringtonine-treated neuron/glia cultures reveals use of novel TISs *in vitro*

To investigate alternative TISs, we used mouse brain-derived primary neuron/glia mixed cultures as our experimental platform because they offer two advantages. First, such cultures can be rapidly and synchronously treated with homoharringtonine (HHT) to arrest initiating ribosomes while allowing elongating ribosomes to run off the transcript^12^. Non-neural cell cultures treated successively with HHT and cycloheximide (CHX), a drug that immobilizes elongating ribosomes^29^ have been used to identify TISs previously^12^. Second, KCl treatment of neuronal cultures induces membrane depolarization, inflow of Ca2+ through L-type Ca+2 channels, and expression of c-Fos and other activity-dependent genes, thus offering a simple yet effective paradigm for studying activity-dependent gene regulation^30–32^.

Accordingly, we utilized neuron/glia cultures with or without HHT and KCl treatments and subjected them to RF following CHX treatment (Fig 1B). Footprint distribution across transcripts revealed a near complete run-off of ribosomes from the proximal 300 nt region of the CDS following 2 minutes of HHT treatment (Fig. 1C). To detect TISs, we focused on this 300 nt region of the most robustly expressed transcripts and developed a set of stringent criteria (see methods) to call high fidelity TISs in both KCl treated and untreated cultures. We detected TISs in 426 transcripts, most of which possessed multiple TISs, with 89.7% having two or more, and only 11.3% possessed a single TIS **(Table S1**, Fig. 1D). Although previous high-throughput studies on TISs have used non-neural cells, we compared our list of alternative TISs to published lists generated from mouse embryonic stem cells^12^ or various human cells^13^ and found that more than 50% of the transcripts in our list overlapped with the lists from the previous studies. These results convinced us that RF on HHT-treated neuronal/glia culture combined with a stringent data analysis pipeline is able to detect alternative TISs in neural transcripts.

Because the number, composition and frame of TISs are important determinants of the proteomic complexity of a cell, we examined these features across the transcripts for which we detected high fidelity TISs (Fig. 1E). On average, a transcript contained 3.18 TISs— with 1.33 uTISs, 0.94 dTIS, and 0.91 aTIS. The aTIS per transcript value of less than 1 resulted from the absence of an initiation peak at the annotated ATG in some transcripts, suggesting that these transcripts might be misannotated and that the dominant protein isoform is not the canonical one in neural cells. ATGs detected at aTISs were always in frame, indicating the robustness of our pipeline and consistent with the presence of ATG as a key contributor to the original annotation. Similarly, near cognate codons such as CTG, TTG and GTG detected at the aTISs were in frame > 99% of the time. In contrast, ATGs and cognate codons detected at uTISs and dTISs were out of frame ≥ 47% of the time. In sum, our data indicate that a brain transcript may use multiple TISs, each at a rate that is regulated in a transcript-specific manner, which can generate multiple protein variants or completely different peptide sequences.

### LC-MS detects protein products of alternative initiation

We next asked if a subset of the alternative TISs results in detectable amount of polypeptides in the neural cells, and were interested to identify which of these were specifically coming from neurons. We noted that most uTISs and dTISs are close to their corresponding aTISs, suggesting that a biochemical assay like Western blot would not clearly resolve the small differences in size between the alternative and normal protein isoforms. Therefore, we examined a quantitative proteome of cultured primary mouse cortical neurons using LC-MS. In spite of the different sensitivities of LC-MS and RF, we clearly detected variant N-terminal peptides resulting from ten of the predicted alternative TISs identified from RF data (Table 1). These included a peptide from an in-frame uTIS lying at the - 63^rd^ nucleotide of *Rtn1*, which encodes reticulon 1, an endoplasmic reticulum protein expressed in neuroendocrine cells and potentially involved in vesicle secretion^33^ and a peptide from an in-frame dTIS lying at the 15^th^ nucleotide of *Uchl1*, which encodes ubiquitin C-terminal hydrolase L1, a neuron-specific deubiquitinating protease with protective roles in neurodegeneration and Alzheimer’s disease^34,35^ (Fig 2 A, B). These results confirm that at least a subset of the TISs detected by our algorithm is utilized in neurons to produce detectable protein and alternative translation does contribute to the diversity in the brain proteome.

**Table 1:**
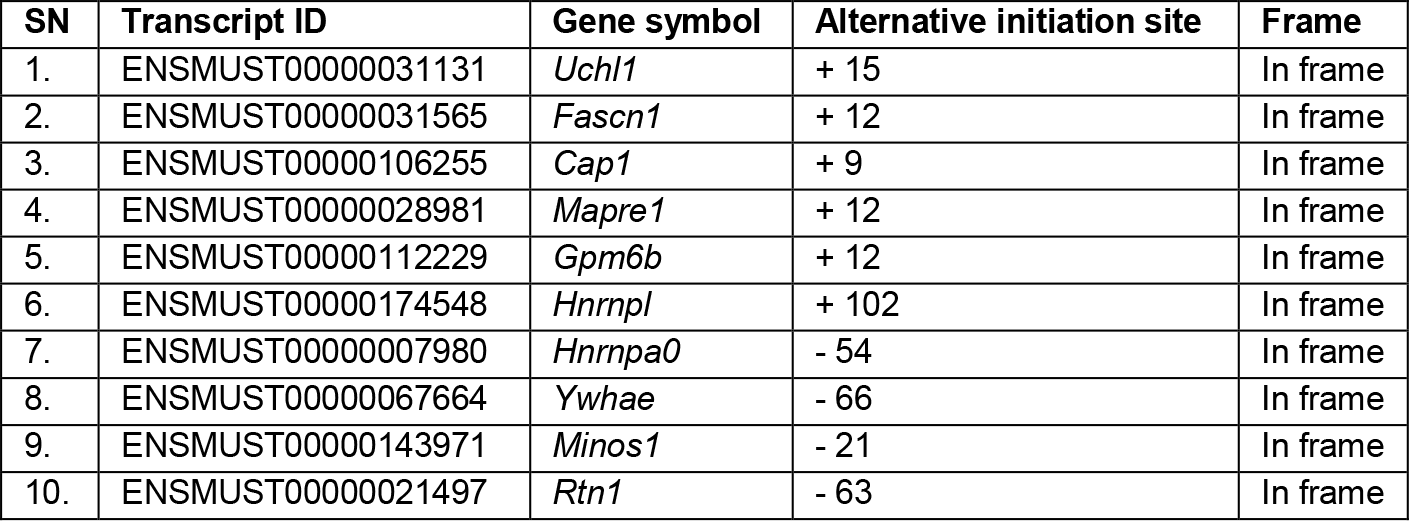
Peptide products of alternative translation initiation sites detected by mass spectrometry analysis of neuronal cultures. “+” and “-” indicate downstream and upstream relative to the canonical initiation site, respectively.

**Figure 2.**
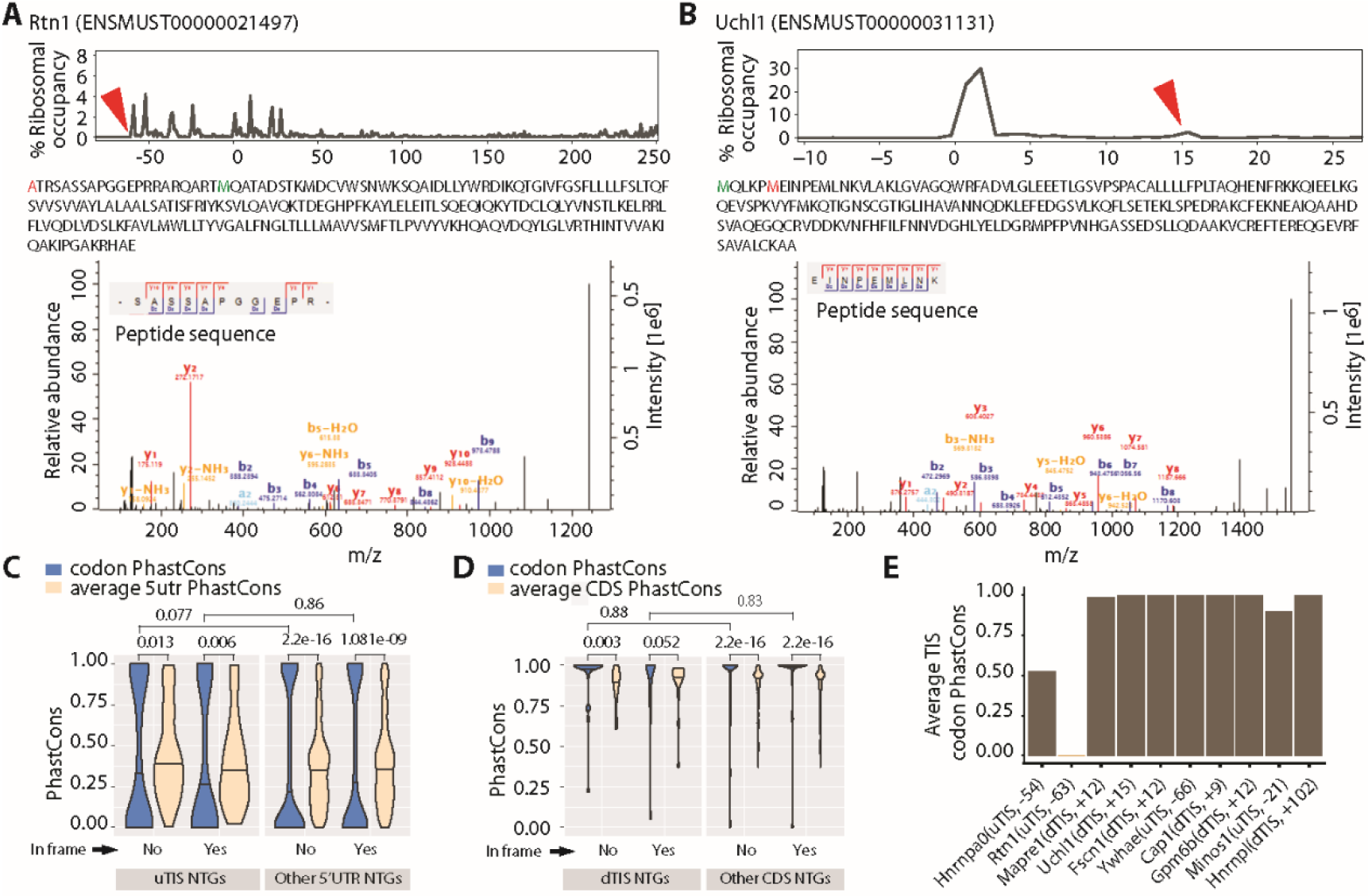
Mass spectrometry detects the peptides resulting from alternative TISs. A-B) Peptides resulting from an in-frame uTIS (*Rtn1*) (A) and an in-frame dTIS (*Uchl1*) (B). Upper panels show the percentage of ribosomes at different TISs (red arrowheads correspond to alternative TISs whose novel products are detected, zeros correspond to aTISs). Middle panels show the peptide sequences with the amino acids corresponding to alternative TISs and aTISs in red and green, respectively. Bottom panels show the tandem mass spectra and mass-to-charge ratios (m/z) of b and y product ions confirming peptide sequences. Alternative TISs for Rtn1 and Uchl1 lie at the −63^rd^ and +15^th^ nts relative to the aTIS, respectively. **C-D)** Conservation of uTISs with respect to 5’UTR (C) and of dTISs with respect to CDS (D). Violin plots show the average of the PhastCons scores for the three nucleotides of the ATG, CTG, GTG or TTG present at the uTISs or dTISs or elsewhere in the 5’UTR or CDS. Wilcoxon test was used to assess the statistical significance. **E)** Conservation of the uTISs and dTISs of mass spectrometry-confirmed peptide products. Plot shows the average of the PhastCons scores of the three nucleotides.

To determine if the identified alternative TISs might represent conserved and thus evolutionarily important adaptations, we next examined whether they were conserved across phyla. We used PhastCons^36^ in a multi-species alignment of 60 vertebrates to compute conservation scores for the NTG codons (ATG, CTG, GTG or TTG) detected as alternative TISs, and tested the hypothesis that these were more conserved than their background sequence (the entire 5’UTRs for uTISs and the entire CDSs for dTISs). We also computed the score for the NTGs not detected as alternative TISs and the respective 5’UTRs or CDSs. Surprisingly, regardless of alternative TISs or not and in-frame or not, 5’UTR NTGs were less conserved whereas CDS NTGs were generally more conserved than their respective backgrounds (Fig. 2C). However, NTGs detected both as uTISs or dTISs were not more conserved than NTGs detected in their respective backgrounds. When we specifically examined the conservation of the ten alternative TISs for which we detected the corresponding peptides with LC-MS (Table 1), we found eight of them to be highly conserved, one only moderately conserved and one not conserved (Fig. 2E). In sum, our analyses suggest that specific alternative TISs, particularly those that are robust enough to generate detectable protein, may be highly conserved. Yet, the significantly *decreased* conservation of NTGs in 5’UTRs, for even many that show initiation, also suggests that some modifications of TISs may be a substrate for evolution of gene regulation across species.

### Transcripts change TISs usage in response to neuronal depolarization

Neuronal activity induces de novo protein synthesis— both from existing mRNAs and via induction of a specific transcriptional program^37–39^. However, it is unclear if it can also alter TIS usage on a given transcript. Therefore, we asked if KCl depolarization of our neuron/glia culture model regulates the use of TISs. For this purpose, we removed 82 transcripts that had only one TIS and/or no aTIS from our list of transcripts. We found that, of the remaining transcripts with at least one alternative TIS, 32.5% exhibited a significant change in the ribosomal occupancies of their aTIS, uTIS or dTIS, with a majority showing increased use of aTIS in response to KCl depolarization **(Fig. S1A)**. More importantly, we observed that an increased use of aTIS was accompanied by a corresponding decreased use of alternative TISs and vice versa, with a few exceptions where changes at both TISs went in the same direction. These results indicate that specific transcripts use their TISs differentially in response to neuronal depolarization.

We next examined the transcripts and directions of effect for this depolarization-regulated use of TISs. The increased use of aTISs by the majority of transcripts was consistent with an increase in de novo protein synthesis from the canonical ORF following stimulation. Transcripts that showed this included *Grina* and *Hspa4*, which code for an NMDA receptor subunit and a heat shock protein involved in the folding of other proteins, respectively^40,41^ **(Fig. S1B,C)**. On the other hand, among the few transcripts that showed an increased use of uTISs included *Kif1b*, which codes for a kinesin superfamily protein involved in transporting synaptic vesicles^42^ **(Fig. S1B,C)**. Interestingly, *Gfap*, which codes for an intermediate filament in astrocytes^43^, also exhibited a modest regulation of its uTIS. Notably, as shown in Fig. S1A, we found that while some of the uTISs and dTISs with altered usage were in-frame, most were out-of-frame, suggesting the regulated production of not only N-terminal protein variants but also novel polypetides following activity. Overall, these findings show that stimulation alters the use of TISs of several transcripts that are involved in diverse functions, indicating a regulated production of alternative protein isoforms in neural cells.

### RF identifies novel C-terminal extensions mediated by stop codon readthrough *in vitro*

Like in frame alternative initiation, stop codon readthrough could enable generation of multiple protein products from a single transcript. As shown in Fig 1A, while uTIS and dTIS can generate protein variants with alternative N-termini, stop codon readthrough can generate a C-terminally extended protein variant. Recently, RF was utilized to identify stop codon readthrough in *Drosophila* embryos^19^. Shortly thereafter, bioinformatics approaches predicted at least 7 readthrough events in human transcripts, of which 4 were validated as capable of >1 % readthrough using luciferase assays in HELA cells^44^. These included *Aqp4* transcript, which encodes a water channel that is highly expressed in astroglia.^45^ A genome-wide screen for transcripts exhibiting readthrough, however, has not yet been conducted in CNS cells.

To profile readthrough events in our neuron/glia cultures, we used the RF dataset from HHT-untreated cultures and designed a pipeline to count the footprints mapping to the 3’UTR of robustly expressed transcripts. We carefully excluded any transcripts where alternative splicing or stop codon single nucleotide polymorphisms might give rise to such footprints (see methods for details). Overall, we identified readthrough of at least 1% in 18 transcripts **(Table S2)**. These include *Aqp4*, as well as malate dehydrogenase 1(*Mdh1*), but not microtubule associated protein 2(*Map2*) (Fig. 3A). To independently validate the ability of these sequences to permit read-through, we cloned a cassette encompassing the distal CDS and the readthrough region of each into a dual luciferase vector so that Renilla luciferase was constitutively expressed while Firefly luciferase was expressed only following stop codon read through^46^ (Fig. 3B). Comparison of the relative luciferase activities confirmed that mouse *Aqp4* sequence permits readthrough of 12.5% when compared to a positive control sequence where the stop codon was mutated to a sense codon (Fig. 3C). This is a level similar to that previously reported for human *Aqp4* in HeLa cells^44^. Likewise, *Mdh1* sequence permits readthrough at 5.5%, while *Map2* sequence is not read through. The addition of an extra stop codon beyond the original stop codons abrogated the readthrough of both *Aqp4* and *Mdh1*, indicating Renilla luciferase expression is not due to a cryptic promoter or internal ribosome entry site sequence. Thus, we have identified specific sequences capable of stop codon readthrough in CNS derived cells.

**Figure 3.**
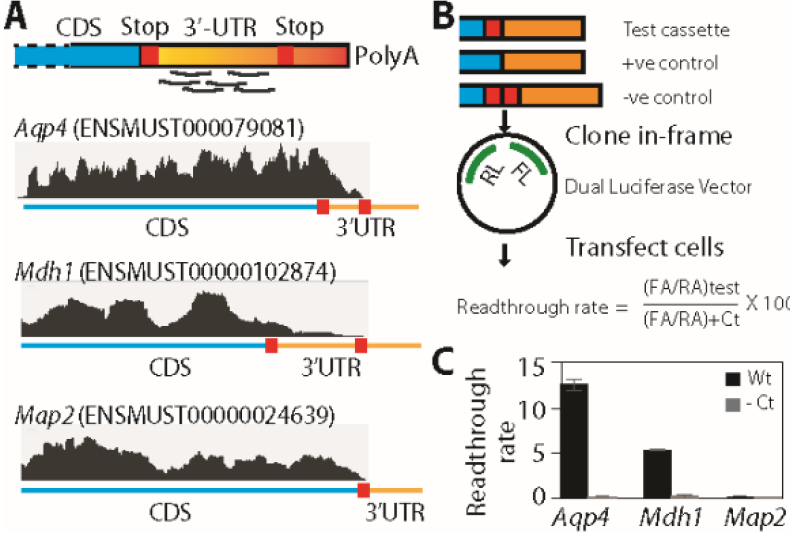
RF of cycloheximide-treated neuron/glia culture reveals stop codon readthrough in vitro. **A)** A schematic of footprint mapping and genome browser tracts with examples of readthrough are shown. *Aqp4* and *Mdh1* display footprints mapping to the readthrough region, whereas *Map2* does not. **B)** Dual luciferase assay assessing the permissiveness of these sequences for readthrough. A cassette spanning the distal CDS and the readthrough region is cloned between the RL and FL such that the latter is expressed in transfected cells only if ribosomes read past the cassette. The stop codon is mutated to a sense codon in the positive control, whereas an extra stop codon is added in the negative control. **C)** Dual luciferase assay shows that *Aqp4* and *Mdh1* undergo readthrough at the rates of 12.5% and 5.5%, respectively, and that Map2 does not undergo readthrough. Readthrough rate is calculated as shown in B (n = 3). CDS, coding sequence; RL, Renilla luciferase; FL, firefly luciferase; RA, Renilla activity; FA, firefly activity.

### TRAP-RF identifies novel, cell type-specific, C-terminal extensions mediated by stop codon readthrough *in vivo*

Although we ruled out several other possibilities that could generate footprints mapping to the 3’UTR and corroborated sequence-mediated readthrough events with luciferase reporters, it was still possible that these events were due to the unnatural conditions of culture. In addition, in culture we cannot definitively establish which cell type is producing the readthrough event. Therefore, we wanted to also assess alternative translation *in vivo*, focusing specifically on readthrough, since, unlike initiation, it does not require pharmacological manipulation in the brain. To this end, we combined RF with TRAP using mouse lines which express Green fluorescent protein (Gfp)-tagged ribosomes in either neurons (*Snap25::Rpl10a-Egfp*) or astrocytes (*Aldh1l1::Rpl10a-Egfp*), thus allowing the purification of neuronal or astrocytic ribosome-bound mRNA^47,48^. Thus, our *in vivo* approach also offered readthrough discovery in a cell type-specific manner.

We subjected the *Snap25::Rpl10a-Egfp* and *Aldh1l1::Rpl10a-Egfp* brains to TRAP followed by RF, an approach we call TRAP-RF (Fig. 4A), similar to an approach recently described by Gonzalez et al^49^. The resulting footprints mapping to the transcriptome were highly reproducible between replicates, indicating a good reliability of the approach we have developed (Fig. 4B, C). We confirmed enrichment of neuronal and astrocyte marker genes in *Snap25::Rpl10a-Egfp* and *Aldh1l1::Rpl10a-Egfp Snap25* TRAP-RF data, respectively, matching prior experiments using these lines^47,48^, indicating we were indeed profiling ribosomes enriched from the cell types of interest (Fig. 4D). We went on and applied the same stringent criteria as used for *in vitro* cultures above to detect readthrough events and identified 50 transcripts with at least 1% readthrough in the TRAP-RF data **(Table S3),** and these were reproducible across replicates. Of these, we found 21 as neuron-enriched, 19 as astrocyte-enriched, and the remaining 10 as non-cell type-specific (Fig. 4E). Finally, we compared the *in vitro* and *in vivo* results and found that 13 out of the 18 *in vitro* candidates had clearly detectable readthrough *in vivo* as well (Fig. 4F), including *Aqp4* and *Mdh1*. For both *in vitro* and *in vivo* readthrough footprints, a metagene analysis revealed peaks at the first and second stop codons, consistent with the fact that translation termination is a slower process and suggesting that the footprints were indeed from ribosomes rather than of other proteins, and that the elongated peptide remained in frame (data not shown). Thus, our TRAP-RF analysis identified both cell type-specific and shared transcripts that might generate C-terminally extended protein variants owing to stop codon readthrough.

**Figure 4.**
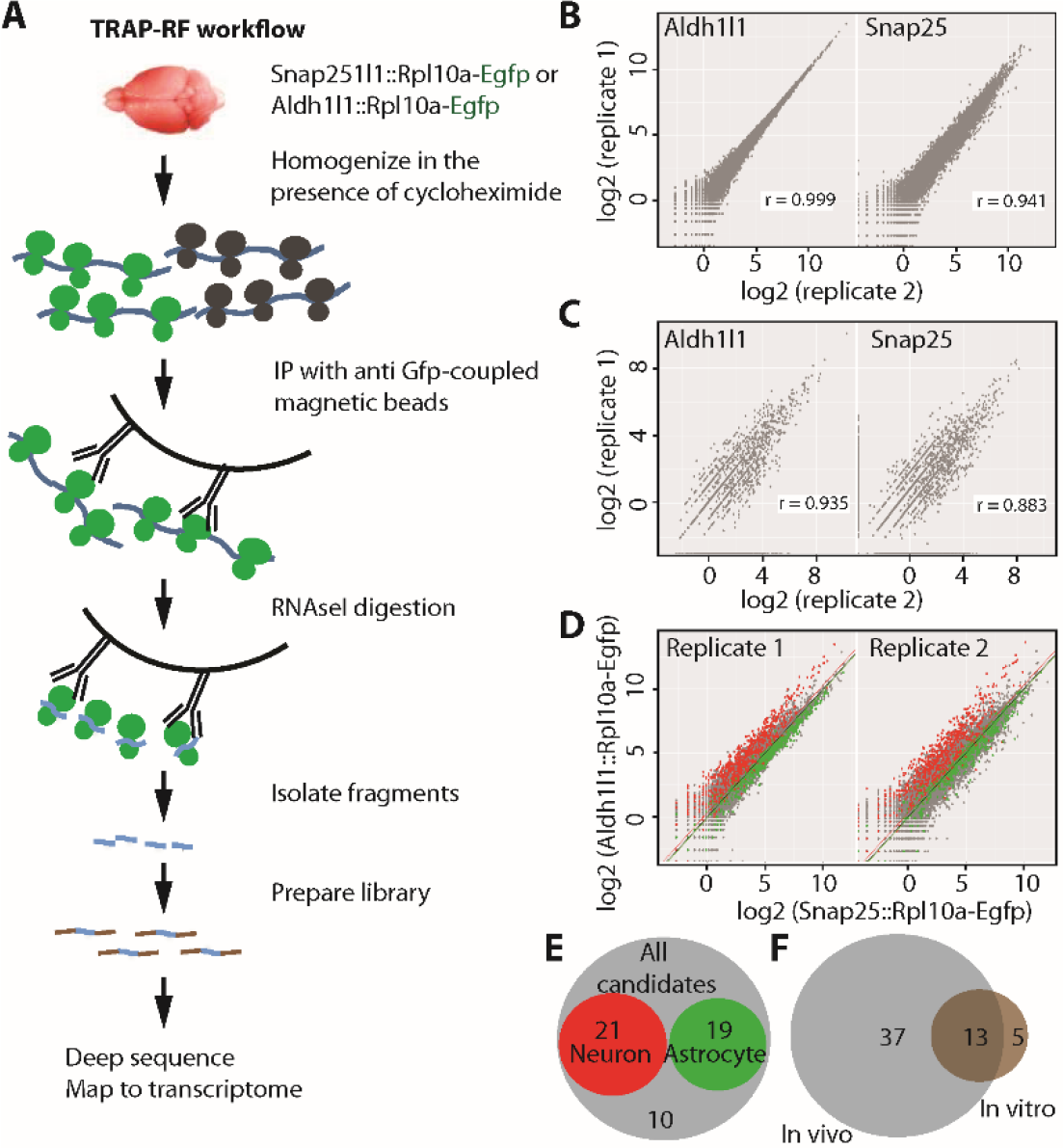
TRAP-RF detects cell type-specific stop codon readthrough in vivo. **A)** TRAP-RF workflow. mRNAs from transgenic brains expressing Gfp-tagged ribosomes in neurons (*Snap25::Rpl10a-Egfp*) or astrocytes (*Aldh1l1::Rpl10a-Egfp*) are affinity-purified with anti Gfp-conjugated magnetic beads, subjected to on-bead digestion with RNAse and the subsequent RF protocol as shown. **B, C)** Log2 (RPKM) of ribosome footprints mapping to the coding sequence (B) and proximal 3’UTR (C) were reproducible between the replicate samples for both *Snap25::Rpl10a-Egfp* and *Aldh1l1::Rpl10a-Egfp*. **D)** Comparison of RF-TRAP samples between neurons and astrocytes shows expected enrichment of neuronal (green) and astrocytic (red) transcripts identified in previous experiments^47^. **E)** Of the 50 transcripts detected undergoing ≥ 1% readthrough in the brain, 21 are neuronal, 19 astrocytic and the remaining 10 non cell type specific. **F)** Of the 18 transcripts with ≥ 1% readthrough in neuron/glia culture, 13 do so in vivo as well.

### Readthrough localizes AQP4 to the perivascular region

We next wanted to test if readthrough is of functional significance. We focused on AQP4, an astrocyte membrane protein shown to have a readthrough version in other species by previous studies^28,44^. First, to determine if a stable readthrough-extended AQP4 protein (designated AQP4X hereafter) is indeed detectable in the mouse brain, we developed a rabbit polyclonal antibody against AQP4X using a peptide from the readthrough region as an epitope. In immunofluorescence staining, the antibody recognized AQP4X but not AQP4 in cell cultures expressing the respective version of AQP4, thus demonstrating sensitivity and specificity **(Fig. S2A, B)**. Western blot on mouse brain lysates revealed that anti-AQP4X recognizes the expected 35 kD band in the adult brain but not in the developing brains of postnatal day 9 or younger mice **(Fig. S3A)**. This is interesting since astrocyte maturation, particularly the endfeet at the blood brain barrier, are not complete until 3 weeks of age^50^. In line with Western blot, immunofluorescence staining showed that AQP4X is abundant throughout the adult mouse brain **(Fig. S3B)**. We noted that AQP4X is significantly concentrated perivascularly, and adjacent to the endothelial cell marker PECAM-1, whereas AQP4 is also abundant farther from the blood vessels **(Fig. S3B, C)**. Similar results were also recently reported by others with an independently raised antibody in rats^28^. Thus, readthrough confers a conserved perivascular localization signal to AQP4.

Because commercial anti-AQP4 antibodies also recognize AQP4X, our immunofluorescence data above could not resolve if it was only AQP4X, or both AQP4X and AQP4, that is localized perivascularly. To address this, we used Adeno-associated virus 9 (AAV9) packaged with cMyc-*Aqp4* (*Aqp4* mRNA with an additional stop codon to prevent readthrough) under the astrocytic promotor Gfap. We reasoned that the cMyc epitope should not be detected adjacent to PECAM-1 if only AQP4 is not perivascular. Immunofluorescence staining of the transduced mouse brains showed that only about 4% of AQP4 is adjacent to PECAM-1**(Fig. S3D, E)**. By comparison, as evident in Fig. S3C, 75% of AQP4X is confined to the perivascular region. Thus, AQP4 mostly lies elsewhere along the astrocyte membrane, whereas AQP4X is mostly perivascular.

AQP4 is expressed in numerous tissues and required for their normal functioning^51,52^. Therefore, we next asked if its readthrough is a brain-specific or a ubiquitous phenomenon. By immunofluorescence staining, we detected AQP4X in the retina, where it was mostly perivascular as in the brain, as well as in the kidneys. However, in contrast to the brain where staining differed from the AQP4 antibodies, in kidney the AQP4 and AQP4X expression patterns were completely overlapping **(Fig. S4)**. Hence, AQP4 readthrough is not a brain-specific phenomenon, however, whether it alters subcellular localization depends on the tissue.

### AQP4 and AQP4X are differentially regulated by gliosis across multiple disease models

Currently it is unclear whether readthrough always occurs at a consistent rate, or if the phenomena is dynamically regulated is response to conditions in the brain. AQP4 overall has been implicated in diverse both physiological and pathophysiological processes in the brain, including brain water homeostasis, edema and ischemia^reviewed in 52^. Given the largely mutually exclusive localizations of AQP4 and AQP4X in the brain, we first examined if they are independently regulated in ischemia. We induced ischemia in adult mice by transient middle cerebral artery occlusion (tMCAO). We chose this approach because in humans, ischemic stroke accounts for nearly 85% of all strokes^53^. In mice, MCAO results in relatively reproducible infarcts in the lateral caudatoputamen and frontoparietal cortex^53^. We performed immunostaining for AQP4 and AQP4X in ischemic mouse brains and quantified their signals in gliotic peri-infarct regions. We found that compared to the regions on the contralesional hemisphere, GFAP-labeled peri-infarct regions showed a 2.5 fold upregulation of AQP4 but only a disproportionate 1.3 fold upregulation of AQP4X (Fig. 5A, B). The differential increase suggested that AQP4 readthrough can be independently regulated from gene expression during pathological processes.

**Figure 5.**
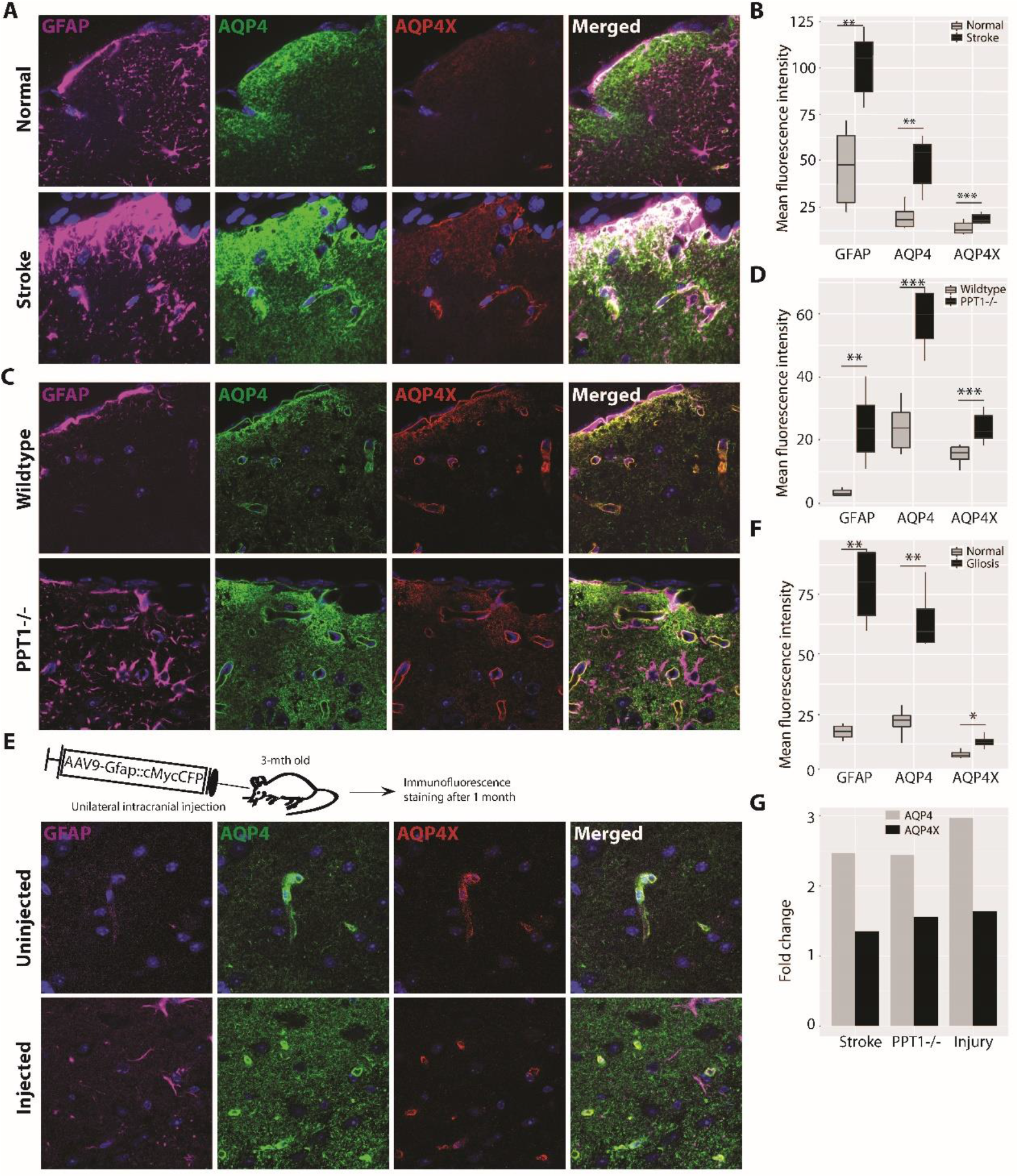
AQP4 and AQP4X are differentially upregulated in gliosis. **A, C, E)** Immunostaining for AQP4, AQP4X and GFAP in brain sections from mice with medial cerebral artery occlusion (A), infantile Batten disease (C) and injury-induced gliosis (E). Normal and gliotic hemispheres of the same sections are imaged in A and E, and age-matched wildtype mice are used in C. **B, D, F)** Fluorescence intensities in A, C and E quantified in B, D and F, respectively, show that AQP4 is significantly more upregulated than AQP4X by gliosis. Regions of interest were drawn to cover the gliotic area and the contralateral normal area in A, whereas whole images were considered in B and C. 3 mice, 6 sections/mouse, paired *t*-test. G) Barplot compiles the relative upregulations of AQP4 and AQP4X from panels B, D and F.

To determine whether this regulation was driven by the ischemia itself, or was more parsimoniously explained as part of gliosis, we next tested if another neurological disease driving reactive gliosis causes a similar differential regulation of AQP4 and AQP4X. For this we used mice lacking palmitoyl protein thioesterase-1 (PPT1), a lysosomal enzyme that is absent in human patients with infantile Batten disease^54^. PPT1−/− mice exhibit several key features of this disease, including neurodegeneration and gliosis^54,55^. We compared the expression levels of AQP4 and AQP4X in PPT1−/− and wildtype brains using immunofluorescence staining and observed that they were indeed differentially regulated in response to gliosis in the knockout brains. While AQP4 expression was 2.5X upregulated, AQP4X was 1.5X upregulated (Fig. 5C,D).

We finally examined if injury-induced gliosis also exerts a similar regulatory effect on AQP4 and AQP4X, simply using tissue from mice having undergone stereotactic injection, which can also induce gliosis. We marked the site of injection using AAV9 expressing cMyc-tagged cyan fluorescent protein (CFP) under the GFAP promoter. A robust expression of cMyc and CFP (data not shown) and mild gliosis were evident along the injected line after one month (Fig. 5E). We then repeated the immunostaining as above and found gliotic regions to express AQP4 at ~ 3 fold and AQP4X at 1.5 fold more intensely as compared to the uninjected contralateral side (Fig. 5E, F). Overall, our results indicate that diseases and conditions accompanied by gliosis differentially upregulate the two isoforms of AQP4 (Fig. 5G).

## Discussion

We present the first study of alternative translation in specific cell types of the CNS, and report that it is a widespread phenomena, with regulated activity and can result in alterations of protein localization and potentially function. Initially identified as viral strategies to diversify protein repertoire^56–58^, non-canonical translational events were later also reported in eukaryotes, including humans^12,18,19,44,59^. Now we have also characterized their extensive use in neural cells. Specifically, we show that hundreds of neural transcripts use uTISs and dTISs *in vitro* and dozens undergo stop codon readthrough both *in vitro* and *in vivo*. Importantly, our MS data demonstrate that several of the uTISs and dTISs do give rise to measurable protein products from neurons. Although MS did not identify any readthrough peptides (more below), we ascertained the presence of AQP4X *in vivo* using a readthrough-specific antibody. Thus, our findings establish that non-canonical translational contributes to the heterogeneity of the neural proteome.

We also provide two lines of evidence that the protein products of non-canonical translation are likely to have functional significance in the brain and are not simply abnormal products destined for immediate degradation. First, we found that they are regulated: more than one hundred transcripts altered their preference for different TISs in response to KCl stimulation. *De novo* protein synthesis in the brain is required for not only housekeeping functions but also specialized functions like long-term potentiation^60^. It is conceivable that the protein isoforms induced by depolarization are involved in carrying out such brain functions. Second, we found that readthrough localizes AQP4 to the perivascular endfeet of astrocytes, and the expression of AQP4 and AQP4X are differentially altered under diverse pathological conditions causing gliosis. Previous studies have reported that cerebral edema, an often fatal complication of stroke^61^, has a better prognosis in *Aqp4* knockout mice^62^. Thus, it may be of clinical significance to determine if AQP4 readthrough suppression ameliorates stroke-associated edema. Studies have also reported that the blood brain barrier is disrupted in infantile Batten disease^63^. Hence, our findings on AQP4X localization and AQP4 and AQP4X regulation may help explain the pathogenesis of this disease. In addition, previous studies have shown that endfeet-polarized AQP4 is required for the efficient removal of amyloid beta from the brain^64,65^, and is lost in Alzheimer’s disease mouse models as well as human patients^66–68^. These findings in concert with ours lead to an intriguing hypothesis: promoting AQP4 readthrough could improve astrocyte-mediated amyloid beta clearance by targeting more AQP4 molecules to endfeet. Thus, future studies identifying regulators of *Aqp4* readthrough could have important implications. Overall, our findings suggest that alternative protein isoforms have implications in both normal brain functions and neurological diseases.

Thus far, we have confirmed alternative isoforms for only a subset of proteins by MS, although RF suggested the possibility of hundreds of such isoforms. To some extent this reflects difference in sensitivity between the approaches, as peptide based measures are inherently more challenging than sequencing based ones such as RF. For instance, Menschaert et al detected only 16 N-terminally extended proteins in mouse embryonic stem cells using MS^16^ although Ingolia et al had shown the possibility of 570 of such proteins in the same cells using RF^12^. In addition, our MS studies were conducted on a more pure neuronal population cultured from a younger age than the RF studies, thus our MS experiment would be insensitive to astrocyte-specific proteins such as AQP4. Thus, negative results from the MS study should be interpreted cautiously. Yet, it is also quite possible that many alternate TISs do not produce stable proteins, but could perhaps still serve regulatory functions. uTISs have been reported to serve regulatory functions in several transcripts^69^, often serving as decoys to suppress the usage of the aTIS in a regulated manner. A prevalence of such sequences would be consistent with the difficulty in detecting protein products, the relatively high frequency of movement of ribosomes from uTIS to aTIS during stimulation, and potentially even significantly lower conservation of NTGs in UTR sequence in general. Evolution across species is posited to be driven more by changes in gene regulation^70^, typically interpreted with regards to enhancer usage^71^, however the same logic may be true at the level of regulation of translation initiation.

Our findings on TIS and readthrough highlight the several potential mechanisms that may warrant future investigation. We found that many alternative TISs show differential ribosomal occupancy in response to KCl treatment, which is known to stimulate neurons by phosphorylating cyclic adenosine 3’,5’-monophosphate response element binding protein (CREB) via an influx of Ca2+ through L-type Ca+2 channels^72^. Previously, we have demonstrated that KCl stimulation of neural cultures modulates the ribosomal occupancy of specific CDSs^39^. Others have shown that KCl-stimulation of neuronal cultures enhances translation initiation by promoting the expression and activity of eukaryotic initiation factor 4E and enhance initiation^73^. What remains to be shown is how exactly the KCl pathway influences the choice of a TIS by ribosomes. It is intriguing to speculate if it does so by regulating cis elements like RNA secondary structures or phosphorylation and regulation of trans factors such as RNA-binding proteins. Likewise, we found that AQP4 readthrough is of functional significance with regard to protein localization and is regulated by gliosis. This, together with the conservation of several of these readthrough peptides to humans^19,28,74,75^, suggests that there are pathways that actively regulate the process. Targeted manipulation of any such pathways may help define the role of stop codon readthrough in the brain or other tissues. Until then, targeting individual transcripts for analysis, as was done for *Aqp4*^28^, may continue to uncover novel roles for individual products arising from such alternative translation events.

## Methods

### Animal Research Committees

All procedures involving mice conformed to the Washington University institutional animal care and use committee.

### Culture

Neuron/glia mix cultures from the cortices of P0 FVB mouse pups were generated as described ^39^. After 7 days *in vitro*, cells were treated with 2 μg/ml HHT ((LKT lab) and compared to parallel cultures treated with DMSO^39^ (Sigma) for 2 m, and then with 100ug/ml CHX (Sigma) for 7 m at 37° C before being lysed for RF^12^. For examining activity-induced TISs, cells were depolarized with 55mM KCl for 3 h^30,31^ before HHT and CHX treatments. RF was conducted in duplicates.

### RF

RF was performed as described^12,39^. Briefly, the cell lysates were treated with RNAseI and subjected to sucrose-cushion ultracentrifugation for pelleting monosomes. mRNA fragments were isolated from the monosomes, size-selected in a polyacrylamide, ligated to cloning linkers, and converted to cDNAs. The cDNAs were then circularized, depleted of rRNAs, PCRed and size-selected and finally deep-sequenced on Illumina Hiseq 2000 (50 bp, single end). Ribosomal profiles of replicate cultures were highly reproducible, with Pearson’s r > 0.96.

### TRAP-RF

P21 *Snap25::eGFP-RpL10a*, *Aldh1l1:: eGFP-RpL10a*, along with their eGFP-negative littermates were euthanized, and their brains frozen in liquid nitrogen and stored at −80° C until use. Two brains were pooled per sample, and replicate experiments were done. TRAP was performed as described^76^ with a few modifications. Briefly, the brains were homogenized in ice in a buffer (20 mM pH 7.4 HEPES, 150 mM KCl, 5 mM MgCl_2_, 0.5 mM dithiothreitol, 100 μg/ml CHX, Turbo DNAse, protease inhibitors, and RNase inhibitors). The lysates were cleared by centrifuging at 2000 xg for 10 m at 4°C and then treated with DHPC (to 30mM, Avanti) and NP-40 (to 1%, Ipgal-ca630, Sigma) for 5 min in ice. Lysates were then further cleared by centrifuging at 20,000 xg for 15 m at 4°C and then mixed with protein L-coated magnetic beads (Invitrogen), previously conjugated with a mix of two monoclonal anti-GFP antibodies^48^, and incubated with rotation for 4 h at 4°C. Beads were washed 5 times with a high-salt buffer (20 mM pH7.4 HEPES, 350 mM KCl, 5 mM MgCl_2_, 1% NP-40, 0.5 mM dithiothreitol, and 100 μg/ml CHX) and then resuspended in normal-salt buffer (150 mM KCl, otherwise as above). To couple to RF, on-bead RNA digestion was performed with RNAse I (Invitrogen) for 1 h with end-to-end rotation, followed by washing three times with normal-salt buffer. Small ribosomal subunits with the mRNA fragments were eluted with ribosome dissociation buffer as described (20 mM pH7.3 Tris-HCl, 250 mM NaCl, 0.5% Triton X-100, 50mM EDTA)^49^. RNA was extracted with phenol-chloroform, quality-tested with Agilent BioAnalyzer, and then subjected to dephosphorylation and subsequent library preparation as described for RF above.

### Analysis of Sequence Data

Culture and TRAP-RF sequencing results were quality-tested using FastQC (version 0.11.2)^77^, and one culture RF sample was removed due to low read depth. Trimmomatic (version 0.32) was used to trim low-quality bases from the ends of reads and to remove adapter sequences^78^. Only fragments 25-35 nt in length were retained for subsequent analysis. Reads aligning to the mouse rRNA were removed using STAR (version 2.3.1z8)^79^. Surviving reads were then aligned, using bowtie2(version 2.2.2)^80^, to the mouse transcriptome (downloaded from Ensembl Release 75) after first removing degenerate sequences from the transcriptome as described^19,81^, retaining both uniquely mapped and multi-mapped reads. Raw and analyzed data are available at GEO:

#### A) TIS identification

For TIS analysis on neuron/glia culture RF data, we focused on the 5’-UTR region and proximal 300bp of the CDS and excluded the downstream region where ribosome runoff after HHT treatment was not complete. In order to calculate TIS score, we first identified ribosomal P sites, which are known to exist at the 12^th^, 13^th^ or 14^th^ nts of the footprints of lengths 28-29, 30-31, and 32-33 nts, respectively ^17,81^. The P site count postions were identified by using the riboSeqR package. To redress small possible errors in the P site estimation, the P site counts were smoothed by taking an average of the P site counts in a 3 nt window, one nt on each side of the initially identified P site. TIS score at every nt position on a transcript was calculated as:

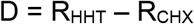

Where R_HHT_ = X_HHT_ / N_HHT_ * 10 and R_CHX_ = X_CHX_ / N_CHX_ * 10. X is the smoothed P site count at a position, and N is the total counts in the 5’UTR and the proximal 300 nt region of the transcript.

The following procedure was used to call TISs:

1. 10 candidate peaks were found in each transcript. The first peak was picked based on highest TIS scores. On each side of this peak, the locations were masked when finding the next peaks if: the location is less than k-bp (k=9) away from the current peak; or TIS scores keep decreasing from the current peak. The next peaks are selected the same way in the unmasked region of the transcript.
2. Assuming that there is at least one TIS in each transcript, the first peak is always called a TIS.
3. If m (m<10) of the 10 candidate peaks were already called as TISs, for the next peak, a score 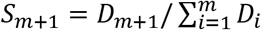 is calcuated, where *D*_*i*_ is the TIS score at the *i*-th peak. If *S*_*m*+1_ > 0.08, this peak is also called as a TIS. Otherwise, only the previous *m* TISs are called for this transcript.

Using this algorithm, TISs were called based on the total P-site counts in KCl treated and untreated samples. TISs for a transcript in KCl+ and KCl− samples were considered the same if they were ≤ 3bp away from each other. For each TIS, the P site count rate was calculated as the P site count over the total counts over all TIS positions. A TIS was excluded if the average rate in KCl+ and KCl− samples was < 0.08. If more than 5 TISs were called for a transcript, only the top 5 TISs with the highest total P site count rates across KCL+ and KCL− samples were kept.

If a NUG codon lay within 3bp away from the called TIS, the TIS was considered overlapping and hence relocated to the position of that NUG codon.

With above criteria, only transcripts with high P-site counts at TIS locations can be used to get reliable TIS calls across samples. Requiring the peak P-sites counts of ≥32 in the considered region of a transcript in the HHT treated group for both KCl+ and KCl−, a total of 426 transcripts were kept for TIS evaluations.

#### B) Readthrough identification

In order to count reads aligning to non-overlapping regions for readthrough analysis in culture and TRAP-RF data, we used the BEDTools intersect command^82^. Then, similar to prior work^19^, we removed reads aligning to the following positions: 12 nts after the start codon; 15 nts before and 9 nts after the first stop codon, and 15 nts after the second stop codon. This resulted in four remaining transcriptomic regions: 5’UTR, CDS, readthrough region, and distal 3’UTR. Reads were counted in each of these regions using the uniq Unix utility.

Readthrough analysis was performed in R. Transcripts were filtered based on CDS counts as follows: in culture RF, transcripts with ≥ 320 CDS counts across all samples were retained, and in TRAP RF, transcripts with ≥ 128 CDS counts across replicates in either the Aldh1l1 or Snap25 samples were retained. Subsequently, to avoid false positive identification of readthrough events because of alternative splicing, if a gene was annotated as having multiple isoforms with stop codons at different genomic locations, only the transcript with the downstream-most stop codon was retained.

Using edgeR^83^, counts were normalized to RPKM, and readthrough rate was defined for each transcript as the ratio of readthrough region RPKM to coding sequence RPKM. Next, for readthrough identification in culture RF and brain TRAP-RF datasets, transcripts with ≥ 32 readthrough region counts across samples were retained. In each dataset, surviving transcripts with ≥ 1% readthrough rate in at least 2 samples were retained for readthrough analysis. Genes with ≥ 100% readthrough or with RPKM(distal 3’UTR) ≥ RPKM(readthrough region), for any transcript isoform or in any sample, were labeled false positives and eliminated.

#### C) Identification of cell type-specific transcripts in TRAP-RF data

For quality assessment and filtering of TRAP-RF data, CDS counts were normalized to CPM using the edgeR package. Due to the expression of some glial transcripts in neuronal samples and vice versa, transcripts were classified as either detectable with high confidence in neurons or astrocytes or as nonspecific background, consistent with prior work^84^. Transcripts were classified as neuron-detectable by first calculating the average log2-fold-change (logFC) in expression between Aldh1l1 and Snap25 samples for genes identified as neuronal markers^85^. Among those transcripts surviving initial CDS count filtering, transcripts with logFC greater than this average plus two standard deviations were identified as detectible in neurons with 96% confidence. Astrocyte-detectable transcripts were identified analogously. Remaining transcripts were designated non cell-type specific.

#### D) Conservation analysis of uTISs and dTISs

For computing conservation scores of uTIS per transcript, phastCons format files were downloaded from UCSC Table Browser, querying the coordinates of uTIS codon from the table of phastCons60way. Next, the average score of the three nucleotides was calculated to define the codon conservation score. Each non-uTIS NTGs in the 5’UTR was annotated using a sliding window approach, and then the codon conservation score was extracted using a similar strategy as for uTISs. These codons were further classified into in-frame and out-of-frame based on their location relative to translation start site. To test uTIS codon score with average score of 5’UTR, average score of the entire 5’UTR given transcripts with uTIS was calculated. As the score distributions appeared bimodal, Wilcoxon’s rank sum test was used to test the pairwise difference.

dTISs were analyzed similarly, except that the corresponding CDSs were considered in place of the 5’UTRs.

### Proteomics

#### A) Sample preparation

Neuronal mass spectrometry (MS) data were obtained similar to the method of Prabakaran et al. 2015^86^. In short, embryonic day 16.5 cortical neurons were cultured for 7 days (3 biological replicates) and depolarized with 55 mM potassium chloride (KCl) for 0, 15, 30, 60, 90, 120 m. For each time point, proteins were extracted in Qproteome mammalian cell lysis buffer (*Complete* mini and *PhosphoSTOP* tablets added (Roche)) using a probe sonicator (15sec, 2x). Protein extracts were reduced with 50 mM dithiothreitol (20 min, 56°C), alkylated with iodoacetamide (30 min, RT), diluted with 8M urea in 50mM triethyl ammonium bicarbonate (TEAB) and digested using filter aided sample preparation (FASP Protein Digestion Kit, Expedeon). Digestion was performed overnight 37°C with 2 ng/μl trypsin (Sequencing Grade Modified Trypsin, Promega) in 50 mM TEAB and peptides were subsequently labeled on-filter using isobaric tags (TMT6plex, Thermo Fisher) according to the manufacturer’s instructions. Labeling was quenched using 5% hydroxylamine and peptides were eluted from the filters using 500 mM NaCl and acidified with 2% formic acid. 10 uL of each peptide eluate were pooled, desalted using C18 silica tips (Nestgroup) to assess labeling efficiency and to adjust sample amounts to ensure 1:1 total protein ratios across channels prior to analysis of the remaining samples. Labeled samples were mixed 1:1 and samples were desalted using Oasis HLB columns (Waters). One half of the sample volume was fractionated using high-pH HPLC fractionation on an Xbridge C18 (3.5μm) 10cm column (Waters) into 55 fractions, the other half by isoelectric focusing on off-gel pH 3-10 Immobiline Dry Strips (GE Healthcare) into 24 fractions. HPLC fractions were combined based on peptide chromatogram intensities into 22 fractions, whereas isoelectric focusing fractions were desalted using C18 silica tips (Nestgroup). Post vacuum drying, fractions were reconstituted in sample buffer (5% formic acid, 5% acetonitrile) for MS analysis.

#### B) MS analysis

Samples were analyzed on a QExactive mass spectrometer (Thermo) coupled to a micro-autosampler AS2 and a nanoflow HPLC pump (Eksigent). Peptides were separated using an in-house packed C18 analytical column (Magic C18 particles, 3 μm, 200 Å, Michrom Bioresource) on a linear 120 min gradient starting from 95% buffer A (0.1% (v/v) formic acid in HPLC-H2O) and 5% buffer B (0.2% (v/v) formic acid in acetonitrile) to 35% buffer B. A full mass spectrum with resolution of 70,000 (at m/z of 200) was acquired in a mass range of 300-1500 m/z (AGC target 3 × 10^6^, maximum injection time 20 ms). The 10 most intense ions were selected for fragmentation via higher-energy c-trap dissociation (HCD, resolution 17,500, AGC target 2 × 10^5^, maximum injection time 250 ms, isolation window 1.6 m/z, normalized collision energy 27%).

Raw data were analyzed by MaxQuant software version 1.5.2.10^77^ and peptide lists were searched against the mouse Uniprot protein sequence database (February 2016, reviewed entries appended with common laboratory contaminants [cRAP database, 247 entries]) appended with the alternative translational products using the Andromeda search engine.^78^ The following settings were applied: trypsin (specificity set as C-terminal to arginine and lysine) with up to two missed cleavages, mass tolerances set to 20 ppm for the first search and 4.5 ppm for the second search. Methionine oxidation and N-terminal acetylation were chosen as dynamic modifications and carbamidomethylation of cysteine and TMT labeling of peptide N-termini and lysine residues were set as static modifications. The minimum peptide length was set to seven amino acids. False discovery rates (FDR) were set to 1% on peptide and protein levels as determined by reverse database search. Peptide identification was performed with an allowed initial precursor mass deviation up to 7 ppm and an allowed fragment mass deviation of 20 ppm. For all other search parameters, the default settings were used.

### Code

Scripts used for sequencing data processing as well as R codes used for downstream analyses are available upon request.

### Dual luciferase assay

*Aqp4* test cassette (the last 15 nts of the CDS + 1^st^ stop + 84 nts before the 2^nd^ stop), a negative control (extra stop added after the 1^st^ stop) or a positive control (1^st^ stop mutated to a sense codon) was cloned between the Renilla and Firefly luciferases of pdLUC, a dual luciferase vector that expresses Renilla luciferase constitutively but Firefly luciferase only if the cloned construct is read by ribosomes^46^. The test and control plasmids were transfected into delayed brain tumor (DBT) glioblastoma cell cultures. After 48 hrs, the two luciferase activities were quantified using the Dual-Luciferase Assay System and GloMax Luminometer with dual injectors (Promega) following manufacturer’s instructions. Readthrough rate was calculated as described ^87^ and as follows:

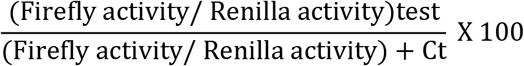

### Immunofluorescence staining

Brain sections, except for those from mice with MCA occlusion, were collected and processed as follows. Adult mice were euthanized and perfused transcardially with 15 ml ice-cold PBS and then 20 ml 4% ice-cold paraformaldehyde in PBS. Brains were harvested, fixed in 4% ice-cold paraformaldehyde overnight, and cryoprotected with 10%, 20%, and 30% ice-cold sucrose in PBS for 4 h, 4 h, and overnight, respectively. Brains where then frozen in OCT (Sakura Inc), sectioned at 40 μm, and stored at 4°C in PBS+0.01%NaN_3_ until staining. For staining, floating brain sections were blocked with 5% normal donkey serum plus 0.3% Triton^®^ X-100 in PBS at room temperature for 1 h, incubated with primary antibody in block at 4°C overnight, washed three times with PBS, incubated with Alexa fluorophore-conjugated secondary antibodies (1:500, Invitrogen) in block at room temperature for 1 h, washed two times with PBS, incubated with 300nM DAPI (Sigma) at room temperature for 5 m, washed two times with PBS, and mounted for confocal imaging (Perkin Elmer).

Brain sections from mice with MCA occlusion were prepared as follows. After transcardial perfusion and overnight fixation as above, the brains were sliced at 1.5 mm, washed with cold PBS for 1 hr and then dehydrated in a series of ethanol concentrations (50%, 70%, 80%, 95%, 100%, 100%, and 100%, each for 1.5 h). After clearing twice with xylene and vacuum-infiltering twice with 58°C paraffin, each time for 1 hr, slices were aligned in molds containing 58C paraffin and embedded in the same paraffin letting the paraffin cool down. Sections were made at 7 um, collected on charged slides, and dried overnight. For staining, slides were first rehydrated by treating in Xylene (3 × 5m, 3 times), 100% ethanol (2 × 10m) times), 95% ethanol (1 × 5m), 80% (1 × 5m), 70% (1 × 5m), 50% (1 × 5m) and distilled water (2 × 5m). Antigen unmasking was then performed by dipping the slides 99°C 10 mM sodim citrate buffer and letting the buffer cool down on bench top for 30m. Slides were then washed 2 × 5m with PBST, and then blocked and stained as described for floating sections above.

Staining of coverslip cultures were done as described previously^39^. Processing and staining of the retinas and kidneys were done as described for floating brain sections above, except that 14 um-thick sections were collected on charged slides.

Primary antibodies and dilutions were: mouse anti-V5 (Sigma, V8012, 1:1000), mouse anti-GFAP (Biogenex, MU020-UC, 1:1000), rabbit anti-cMyc (Sigma, C3956, 1:100), mouse anti-cMyc (Santa Cruz, 9E10, 1:100), goat anti-AQP4 (Santa Cruz, SC-9888, 1:100), rat anti-PECAM-1 (BD Pharmigen, 550274, 1:50), and rabbit anti-AQP4X (made in collaboration with Cell Signaling Technology,1:1000). The anti-AQP4X was a polyclonal antibody generated by immunizing rabbits with a synthetic peptide corresponding to residues surrounding Asp333 of mouse AQPX and purifying the antibody with protein A and peptide affinity chromatography.

### Western blot

Mice were euthanized, and their brains were rapidly homogenized on ice in RIPA buffer (50 mM pH 8.0 Tris-HCl, 150 mM NaCl, 1% NP-40, 0.5 % sodium deoxycholate, 0.1% SDS, 1mM NaVO4, 1mM NaF, Roche complete protease inhibitor tablet). Lysates were cleared by centrifuging at 16,000 xg at 4°C for 10 m. Protein concentration was measured using a BCA kit (ThermoScientific). An aliquot of 30 μg of protein was boiled for 5 m in an equal volume of 2x Laemmli buffer (BioRad) and electrophoresed in Mini-Protean Precast gels (BioRad) for 1h at 120 V. Proteins were transferred to a polyvinyl membrane using Semi-Dry Transfer Cell (BioRad) for 30 m at 10 V and following manufacturer’s instructions. The blot was then blocked in 5% non-fat dried milk in tris-buffered saline with 0.1 % Tween-20 (TBST) for 1 h at room temperature, incubated with anti-AQP4X in block (1:2000, Cell Signaling Technology) overnight at 4°C, washed three times with TBST, incubated with HRP-conjugated secondary antibody in block (1:5000, BioRad) for 1 h at room temperature, washed three times with TBST, and finally developed using Signal-Fire ECL reagent (Cell Signaling Technology) and My ECL imager (ThermoScientific).

### Transfection

Delayed brain tumor cell line was grown in 6 well plates in DMEM with 10 fetal bovine serum (Sigma) and 1x penicillin and streptomycin. At 60% confluency, the cells were transfected with 1.5 μg of plasmids using Lipofectamine 2000 (Invitrogen) and following manufacturer’s instructions. Medium was changed 24 h post-transfection, and immunofluorescence staining was done 48 h post-transfection.

### AAV preparation and intracranial injection

The *Aqp4* cDNA with a 5’ cMyc tag and an extra stop codon beyond the first stop codon and CFP cDNA with a 5’ cMyc tag were cloned into pAAV-GFAP-EGFP after removing the EGFP with AgeI and HindIII. The vectors were packaged into AAV9 by the Hope Center Viral Vector Core at Washington University. Two uL of the viruses were intracranially injected into P1 (AAV9-GFAP-cMyc-Aqp4) or P90 (for AAV9-GFAP-cMyc-CFP) mice. After 3 weeks, the mice were processed for floating brain section immunofluorescence as described above.

### tMCAO

tMCAO was performed as described previously^88^. Briefly, adult C57Bl/6 mice were anesthetized with isoflurane, and the left common carotid artery (CCA) was exposed through a midline cervical incision. A 6.0-gauge nylon suture coated with silicone was inserted in the CCA lumen and advanced through the internal carotid artery to the origin of the MCA. Interruption of the blood flow in the MCA territory was confirmed with laser Doppler. After 60 m, the suture was withdrawn, and the reperfusion of the MCA territory was confirmed by inspection with an operating microscope and more distally by laser Doppler. After the procedure, animals recovered in an incubator before returning to home cages. The brains were harvested after 24 h and processed for immunofluorescence staining as mentioned above.

## Acknowledgements

We thank Gary Loughran for the dual luciferase plasmid, the Hope Center Viral Vectors Core at Washington University in St. Louis for viral vector preparation, Mike Vasek for assistance with image analysis, and the members of the JDD laboratory for scientific discussions. This work was supported by the NIH (1R21DA038458, 1R01NS102272), the Hope Center, and the Children’s Discovery Institute of Washington University in St. Louis to JDD, the McDonnell Center for Cellular and Molecular Neurobiology Postdoctoral Fellowship to DS, the NIH R01GM112007R to JAJS, R01NS084028 to JML, R01NS043205 to MSS and P30CA91842 and UL1TR000448 to the Genome Technology Resource Center at Washington University in St. Louis. JDD is a NARSAD independent investigator of the Brain and Behavior Research Foundation. JDD has received royalties related to TRAP in the past.

